# Genome Editing With Targeted Deaminases

**DOI:** 10.1101/066597

**Authors:** Luhan Yang, Adrian W. Briggs, Wei Leong Chew, Prashant Mali, Marc Guell, John Aach, Daniel Bryan Goodman, David Cox, Yinan Kan, Emal Lesha, Venkataramanan Soundararajan, Feng Zhang, George Church

## Abstract

Precise genetic modifications are essential for biomedical research and gene therapy. Yet, traditional homology-directed genome editing is limited by the requirements for DNA cleavage, donor DNA template and the endogenous DNA break-repair machinery. Here we present programmable cytidine deaminases that enable site-specific cytidine to thymidine (C-to-T) genomic edits without the need for DNA cleavage. Our targeted deaminases are efficient and specific in *Escherichia coli*, converting a genomic C-to-T with 13% efficiency and 95% accuracy. Edited cells do not harbor unintended genomic abnormalities. These novel enzymes also function in human cells, leading to a site-specific C-to-T transition in 2.5% of cells with reduced toxicity compared with zinc-finger nucleases. Targeted deaminases therefore represent a platform for safer and effective genome editing in prokaryotes and eukaryotes, especially in systems where DSBs are toxic, such as human stem cells and repetitive elements targeting.

---

Genetic modification of mammalian cells has been greatly facilitated by the development of customized zinc finger (ZF)^2, 3^-, transcription activator-like effectors (TALEs)-^1^ nucleases^4, 5^ and clustered regularly interspaced short palindromic repeat/Cas9 (CRISPR/Cas9^6–8^). These programmable nucleases create targeted genomic DNA breaks that enhance the incorporation of exogenous DNA sequences via homology-dependent repair (HDR). However, HDR faces numerous limitations: First, the nuclease-induced DSBs are associated with genomic aberrations and cytotoxicity^9^, which is further compounded when targeting multiple genomic loci^10^. Second, HDR is highly inefficient compared to the competing non-homologous end joining (NHEJ) pathway^11, 12^, which results in generation of undesired mutations instead of the intended genetic modifications. Third, in vivo genome editing remain challenging due to the difficulty in delivering donor DNA at sufficient concentration and the diminished HDR efficiency in somatic cells^13^. To bypass these limitations, we developed targeted deaminases, multiplexable genome-editing tools that work independently of DNA cleavages and donor DNA requirements. We further show that our platform alleviates the toxicities associated with traditional nuclease-mediated genome editing.

Deaminases are naturally occurring proteins that operate in various important cellular processes. Activation induced deaminase (AID) and apolipoprotein B mRNA editing enzyme catalytic polypeptide-like family proteins (APOBECs)^14^ are cytidine deaminases critical to antibody diversification and innate immunity against retroviruses^14^. These enzymes convert cytidines to uracils in DNA. If DNA replication occurs before uracil repair, the replication machinery will treat the uracil as thymine, leading to a C:G to T:A base pair conversion^15^. This elegant editing mechanism suggests a simple and effective genome editing approach that circumvents the limitations associated with nuclease-based approaches.

## Design of targeted deaminases

We first engineered targeted deaminases by fusing each candidate deaminases (APOBEC1, APOBEC3F, APOBEC3G (2K3A)^16^ and AID) with a sequence-specific ZF (recognizing the 9bp DNA sequence 5’-GCCGCAGTG-3’^17^ (Fig. 1a). Based on available structures^18^, we inferred that the deaminase catalytic domains reside at the C-terminus. Therefore, to prevent steric hindrance to catalysis, we tethered the ZF to the N-terminus of the deaminases, separated by a four amino-acid linker (Fig. 1c). To determine editing efficiency *in vivo*, we integrated a single-copy GFP reporter into the *E. coli* genome^19^ (Fig. 1b and Supplementary Methods 2) in which the GFP is normally not expressed due to a ‘broken’ start codon (‘ACG’). Correction of the genomic ACG to ATG by targeted deamination would restore GFP protein expression, thereby producing GFP-positive cells quantifiable by flow cytometry. Among the four chimeric deaminases we tested, ZF-AID induced comparatively more robust correction efficiency (Fig. 1c). We confirmed the intended ACG-to-ATG conversion in 20/20 randomly chosen GFP+ bacterial colonies, as assessed by Sanger sequencing. Therefore, ZF-AID introduces C→T mutations at the locus specified by the fused DNA-binding module.

**Figure 1.**
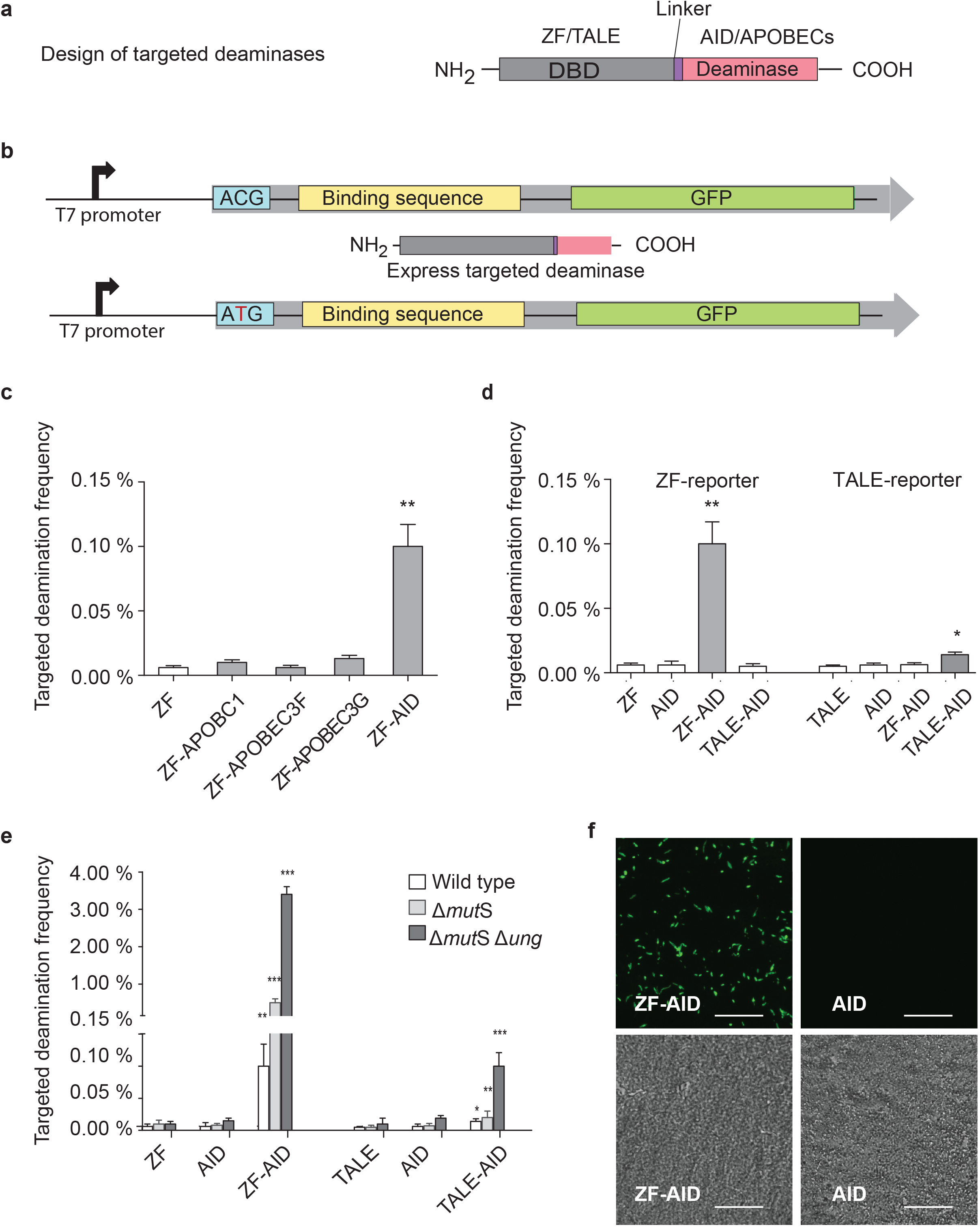
Design and targeted deaminase activity of chimeric deaminases in *E.coli*. **a**, Schematic representation of the design of targeted deaminases. The DNA binding domain (DBD), either ZF or TALE, was fused to N-terminus of the deaminase with a certain linker. **b**, Experimental overview: we integrated a GFP cassette (top) consisting of a broken start codon ACG, DNA binding sequence, and the GFP coding sequence into the bacterial genome. We subsequently transformed targeted deaminases (middle) in pTrc-kan plasmid (**Supplementary Method1**) into the strain and induced protein expression. Targeted deamination of the C in the broken start codon leads to a ACG→ATG transition (bottom), rescuing GFP translation which is quantifiable via flow cytometry. **c**, ZF-deaminases were tested for targeted deaminase activity by measuring GFP rescue. ZF, ZF-APOBECs (ZF-APOBEC1, ZF-APOBEC3F, ZF-APOBEC3G) or ZF-AID indicate cells transformed with plasmids that express ZF, ZF-APOBECs or ZF-AID respectively. All error bars indicate s.d. (All t-tests compare ZF-deaminases against the ZF control. P_value_ < 0.05 *, P_value_ < 0.01 **, P_value_ < 0.001 ***, n=4). **d**, GFP rescue by ZF-AID and TALE-AID in the ZF-reporter and TALE-reporter strains.(All *t*-tests compare the fusion deaminases against the AID control. P_value_ < 0.05 *, P_value_ < 0.01 **, P_value_ < 0.001 ***, n=4). **e**, GFP rescue by ZF-AIDs and TALE-AID in (wild type), (Δ*ung*), and (Δ*mutS* Δ*ung*) strains. All error bars indicate s.d.. (All t-tests compare the fusion deaminases against the AID control. P_value_ < 0.05 *, P_value_ < 0.01 **, P_value_ < 0.001 ***, n=4). **f**, *E.coli* (Δ*mutS* Δ*ung*) cells imaged under fluorescence(upper) and phase contrast(lower) after expression of ZF-AID or AID for 10 hours. Top, Scale bar: 20μm. More detailed structures and sequences of the fusion proteins and reporters are shown in Figure 2a, 2c, **Supplementary figure 4** and **Supplementary Sequence**.

Our targeted deaminases are modular, such that the DNA-addressing component is interchangable with unrelated DNA-binding domains. To demonstrate this, we developed a TALE-AID fusion (recognizing a different binding sequence 5’-TCACGATTCTTCCC-3’^20^) corresponding reporter *E. coli* strain (Fig. 2c). Induction of TALE-AID for 10 hours led to GFP expression in 0.02% of the reporter population, lower than that from ZF-AID, but nonetheless significantly higher than with TALE or AID expression alone (t-test, two-tailed, P_(TALE-AID, TALE)_ =0.0069, P_(TALE-AId, AID)_ =0.0186; n=4) (Fig. 1d). Importantly, GFP expression is dependent on correct sequence recognition, because TALE-AID and ZF-AID do not induce GFP expression in reporter E. coli cells lacking the cognate target sequences (Fig. 1d). Additional target DNA sequences do not increase editing efficiency, suggesting that a single ZF-AID or TALE-AID is sufficient for editing (Supplementary Data 1 and Supplementary Fig. 1). Thus, ZF-AID and TALE-AID converts C-to-T at sequence-defined genomic loci.

**Figure 2.**
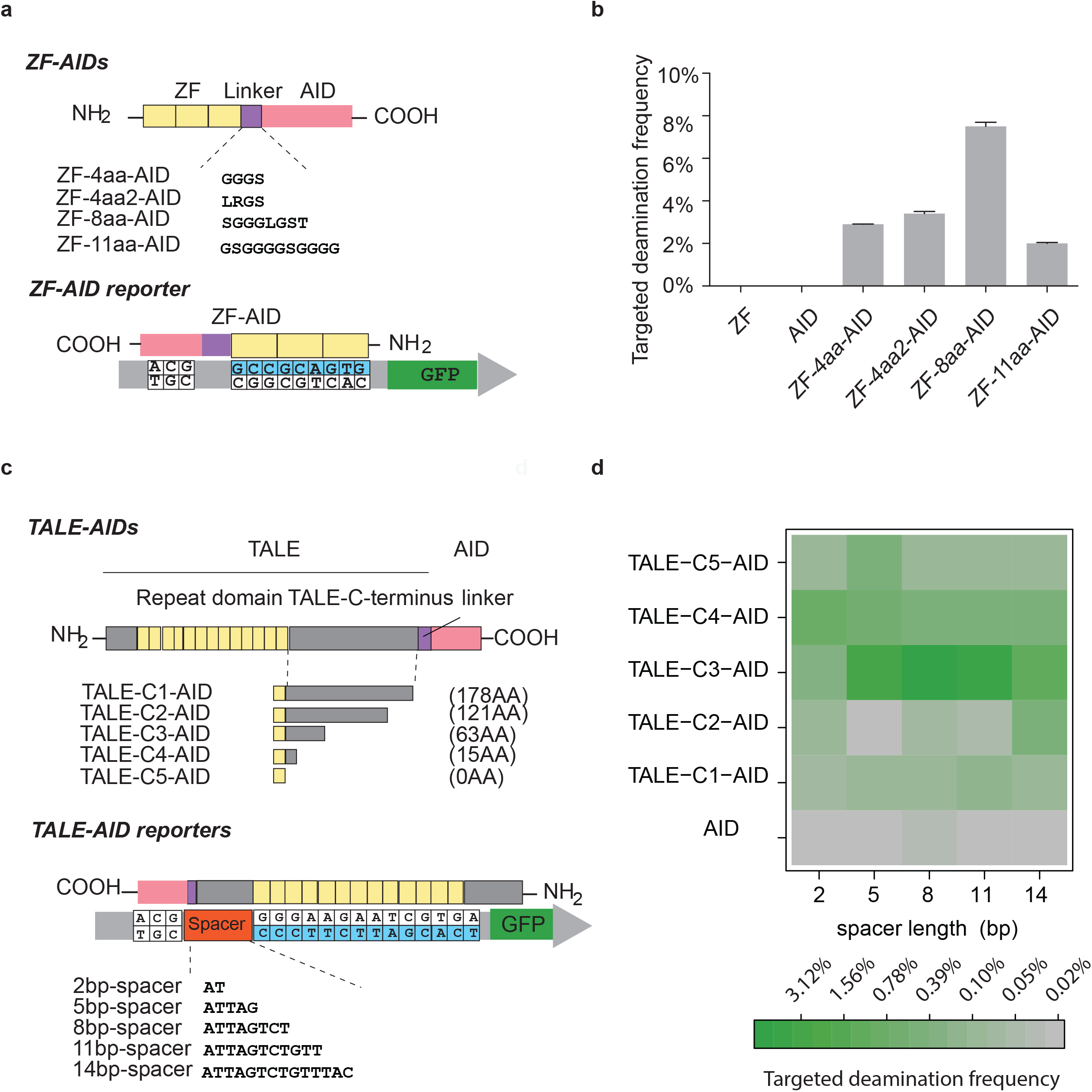
Optimization of targeted deamination frequency of AID fusions in *E.coli*. **a**, Schematic representation of ZF-AIDs variants tested for targeted deaminase activity (upper) and the reporter (lower) with the ZF-recognition sequence in blue. **b**, GFP rescue by expression of the four ZF-AIDs variants and ZF or AID domains alone. All error bars indicate s.d.. **c**, Schematic representation of TALE-AIDs and the reporters tested for targeted deaminase activity. Five TALE-AIDs (upper) with different TALE C-terminus truncations (C1 to C5) were constructed, with the remaining C-terminus lengths shown in parentheses. Full TALE-AID protein sequences can be found in **Supplementary Sequence 2**. Five reporters were constructed (lower) with different spacer lengths (2bp, 5bp, 8bp, 11bp) between the broken start codon and TALE DNA binding motif. The TALE binding site on the GFP reporter is shown in blue; the TALE N-terminus segment specifies the 5′ thymine base of the binding site. **d**. All five TALE-AIDs were tested for targeted deaminase activity on all five reporters (**2c**). Green and grey encode high and low GFP rescue, respectively.

The results demonstrated feasibility of using targeted deaminases for genome editing, but editing efficiency was low. We reasoned that the endogenous uracil repair pathways could reverse the targeted deamination, which would limit the desired C-to-T conversion. Therefore, we knocked out *mutS* and *ung* (Supplementary Method 2), two genes critical for uracil repair. Editing by ZF-AID increased to 0.5% (5-fold) in the *ΔmutS* knockout, and to 3.5% (35-fold) in the *ΔmutS Δung* double knockout (Fig. 1e). Similarly, editing by TALE-AID increased to 0.1% (7-fold increase) in the *ΔmutS Δung* knockout (Fig. 1e). We confirmed the GFP fluorescence signal by microscopy (Fig. 1f) and confirmed the C:G →T:A transitions by sequencing the *gfp* gene of 20 randomly chosen GFP+ colonies from both the ZF-AID- and TALE-AID-induced population. Hence, suppression of uracil repair increases editing frequencies from the targeted deaminases. All subsequent experiments in *E.coli* were done in the *ΔmutS Δung* background.

## Optimization of targeted deaminases

We next conducted structural optimization of the targeted deaminases by varying linker lengths and sequence compositions^21, 22^ (Fig. 2a). While tested variants all led to robust GFP rescue, a longer linker length improved editing frequencies, with ZF-8-aa-AID achieving 7.5% GFP+ frequency after 10 hours (Fig. 2b) and 13% after 30 hours of induction (Supplementary Fig. 2a). Sequence composition of the linker also influences editing frequencies (t-test, two tailed, p=0.0032, n=4). Hence, the linker determines performance of the overall construct.

Our initial TALE –AID (hereafter referred to as TALE-C1-AID) is less efficient than the ZF-AIDs (Fig. 1e). Given the importance of the linker between the DNA-binding module with the deaminase, we proceeded to investigate if truncation of the 178aa^20^. C-terminus could increase TALE-AID activity (Fig. 2c). Truncations were chosen at *in silico* predicted loop regions. We also constructed five bacterial GFP reporter strains, each with a genomic *gfp* locus carrying a broken start codon 2, 5, 8, 11, or 14 bp upstream of the TALE binding site (Fig. 2c). Targeted deamination frequencies were then measured by GFP rescue frequency and compared in a 5-by-5 matrix of TALE-AIDs and reporters (Fig. 2d). TALE-AID truncations showed significantly higher GFP rescue over that of TALE-C1-AID (Fig. 2d), with TALE-C3-AID achieving a genomic editing frequency of 2.5% on the 8bp-spacer reporter after 10 hours of induction (Fig. 2d), and 8% following 20 hours of induction (Supplementary Fig. 2b). Interestingly, TALE-C3-AID outperformed all other constructs regardless of the reporter spacer length, suggesting that this chimeric protein has an intrinsically optimal structure out of the TALE-AIDs tested. These results for ZF-AIDs and TALE-AIDs thus reveal important design considerations for engineering efficient targeted deaminases (Fig 2e).

## Specificity of targeted deaminases

Having investigated and improved deaminase targeting frequency, we next characterized targeting specificity using the following three methods: 1) investigating the effect of point-mutations in the target DNA sequence; 2) deep-sequencing the GFP locus of the population; and 3) whole-genome sequencing of three GFP+ clones.

For the first assay, we show that single-nucleotide change in the cognate target sequence led to 4–8 fold decrease in observed editing rates (Fig. 3a), indicating that ZFP-8aa-AID is specific to the target locus. We next investigated the specificity of TALE-AID by individually varying each nucleotide in the TALE recognition site to the second most preferred base^1^ for that position (Fig. 3b). Interestingly, TALE-C3-AID, which was designed to recognize a 14bp sequence, showed strong sequence specificity only for the first 8bp proximal to the target site (5’ TTCTTCCC 3’ in the TALE recognition site). For reasons that remain to be investigated, sequence alterations at more distal positions in the TALE binding site led to variable targeting frequency (Fig. 3b).

**Figure 3.**
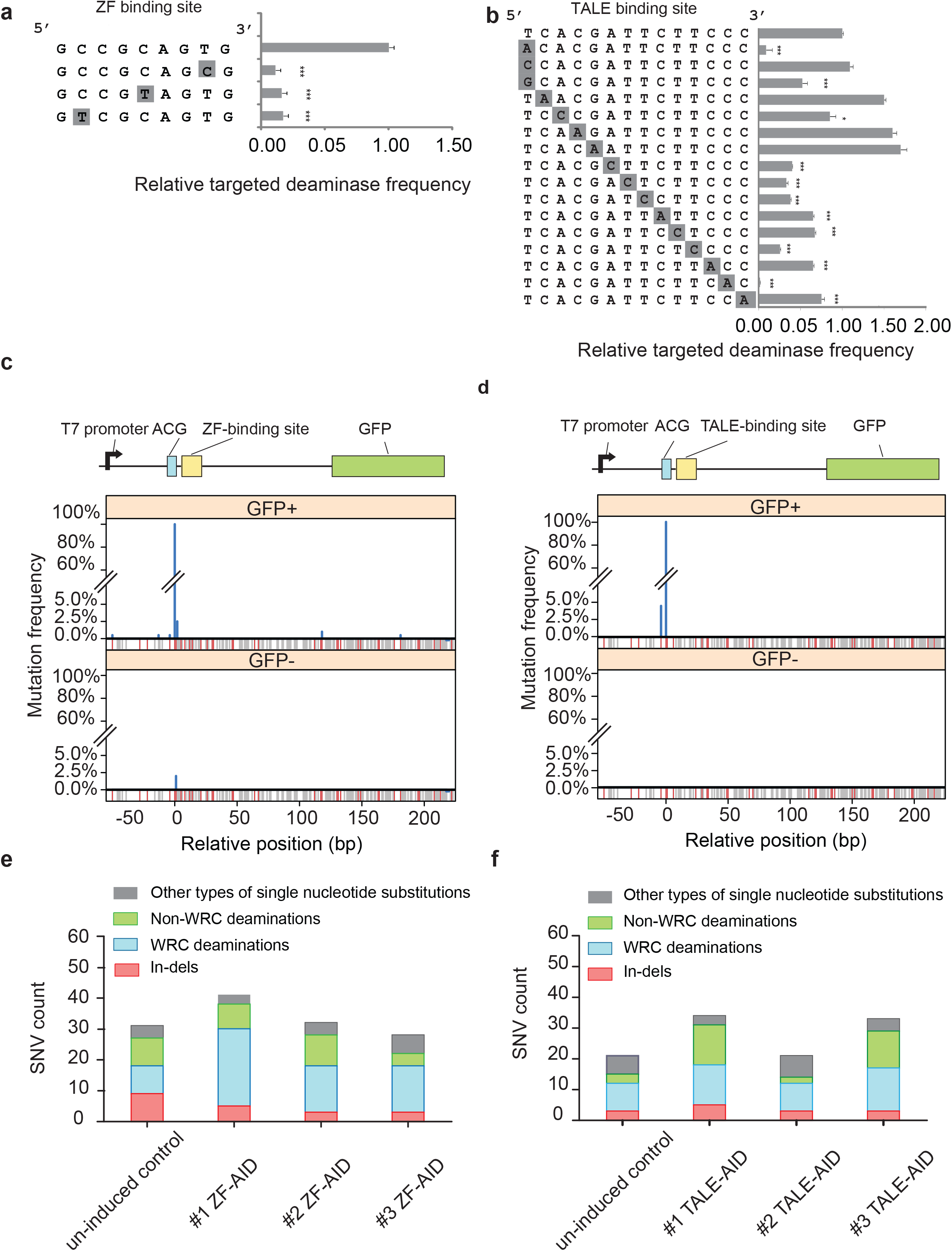
Test of the specificity of AID fusions. **a**, Test of ZF-8aa-AID sequence specificity using a GFP reporter with point-mutated ZF binding sequences. *t*-tests compare each mutated site against the unmodified site (top). P_value_ < 0.05 *, P_value_ < 0.01 **, P_value_ < 0.001 ***, n=4. All error bars indicate s.d.. **b**, Test of TALE-C1-AID sequence specificity using a GFP reporter with point-mutated TALE binding sites. *t*-tests compare each mutated site against the unmodified site (top). Pvalue < 0.05 *, Pvalue < 0.01 **, Pvalue < 0.001 ***, n=4. All error bars indicate s.d.. Note that we altered the first nucleotide, a TALE-N terminus-specified thymine, to other three nucleotides individually, while we changed other nucleotides in the TALE recognition domain to the nucleotide mostly likely to be recognized^1^. **c**. Mutation location and spectrum in the GFP gene of GFP+ and GFP-cells collected after ZF-8aa-AID induction. A schematic structure of the GFP gene is shown above the mutation frequency along the gene’s length among 200 Sanger sequenced colonies of each cell population. Gray lines indicate positions of C/G nucleotides; red lines indicate occurrences of the AID preferred motif (WRC). **d**. Mutation spectrum on the GFP gene of GFP+ and GFP− cells collected after TALE-C1-AID induction. **e**, Whole-genome SNV profiles of strains with/without ZF-AID induction. SNVs that may stem from cytosine deamination (C/G→T/A) are in either green (if C was in the AID-preferred WRC motif) or blue (all other Cs) bars. **f**, Whole-genome SNVs profiles of strains with/without TALE-AID induction. Color schematic is the same as 3e.

Next, to examine on-target editing at single-bp resolution, we sorted 10,000 GFP+ and 10,000 GFP-cells after 30 hours of ZF-8aa-AID induction, and randomly isolated 200 individual colonies from each population. We Sanger sequenced 1kb surrounding the *gfp* target site and, as control, the constitutively expressed *gapA* gene, which lies 1.9Mbp away from *gfp*. In the GFP+ population, all colonies harbored the intended C→T transition in the *gfp* start codon. Interestingly, 5.5% (11/ 200 colonies) of these colonies carried additional C→T transitions in the GFP transgene (Fig. 3c). Most of these additional mutations were confined in a +/− 15bp region flanking the ZF binding site, mutations >150bp away were also detected, suggesting catalytically processivity of AID^23^. In the GFP-population, the only variant detected over 200 colonies was a G□A transition 1bp away from the intended target site (ACG□ACA) that is present in 2% of the population (Fig. 3c). No mutations were found in *gapA* in any colony from the two populations. We next repeated our assay using TALE-C3-AID. In the GFP+ population, besides the intended C→T mutation, an additional C→T mutation 4bp upstream of the intended site was found in 4.5% population (9/ 200 colonies, Fig. 3d). No other off-target mutations were detected in the GFP coding sequence or in the GFP-cells. We concluded that targeted deaminases have residual processivity that mutate nearby C’s within a +/-15bp window of the target DNA sequence.

Finally, to assess genome-wide off-targeting, we sequenced with ~50X coverage the genomes of three GFP+ colonies edited by ZF-8aa-AID, and three colonies edited by TALE-C1-AID, and compared them to control GFP-colonies in which the expression of deaminases had not been induced. We did not observe increased indel mutations in the ZF/TALE-AID expressing clones (Fig. 3e and 3f, Wilcoxon, P_value_=0.7109). However, we detected elevated levels of genome-wide C:G→T:A transitions in WRC sequence motifs following expression of targeted deaminases (Wilcoxon, P_value_=0.02, t-test) (Fig. 3e and 3f). In addition, we did not find any off-target mutations at predicted ZF/TALE off-target sites. The fact that off-target mutations are enriched at WRC motifs - the canonical AID recognition sequence^24^-suggest that off-target mutations derived from intrinsic activity of AID.

## Human genome engineering using targeted deaminases

Given the intense interest in precise genomic editing for human biomedical studies, we tested functionality of our targeted deaminases in human cells. We constructed a mammalian reporter in which an EF1α promoter drives expression of a GFP harboring a broken-start-codon (ACG), followed by an IRES-mCherry selection marker. The reporter construct was stably integrated into HEK293FT cells by lentiviral transduction and a clonal cell line was isolated by FACS sorting (Fig. 4a). The optimized ZF-AID construct (ZF-8aa-AID) was then delivered into the reporter cell line via transfection (Fig. 4a). Following 48 hours of ZF-8aa-AID expression, 0.12% of transfected cells turned GFP+. We next constructed ZF-AID^ΔNES^ by truncating the 15aa from the C-terminus of AID, which contains a strong nuclear export signal^25^ and regions that interact with mismatch repair proteins^26^. This is expected to correctly localize the ZF-AID to the nucleus and decouple AID from the mismatch repair pathway. The expression of ZF-AID^ΔNES^ significantly increased GFP+ cell frequency compared to full-length ZF-AID (Fig. 4b) (0.56%, t-test, two-tailed, n=4, Pvalue=0.0013). Encouraged by our *E.coli* study, we examined if inhibiting the counteracting pathways of uracil repair and mismatch repair would increase C-to-T transition in human cells. Interestingly, the combination of the UNG inhibitor UGI^27^ and MSH2 shRNA increased ZF-AID^ΔNES^– mediated editing efficiency to 2.5% (Fig. 4b). In contrast, the expression of ZF_GFPINI_-Aid^δnes^, a fusion protein whose zinc finger domain targets a site 265bp away from the GFP start codon, resulted in minimal GFP rescue (Fig. 4a,c), suggesting that genome editing by ZF-AID^ΔNES^ is sequence-specific. Successful C:G→T:A targeting of the broken start codon was confirmed by Sanger sequencing of the *GFP* locus in 8/8 stable GFP+ colonies. Therefore, engineered deaminases are capable of efficient sequence-specific genome editing in HEK293FT cells, and editing efficiency can be significantly increased by inhibiting the uracil repair pathway.

**Figure 4.**
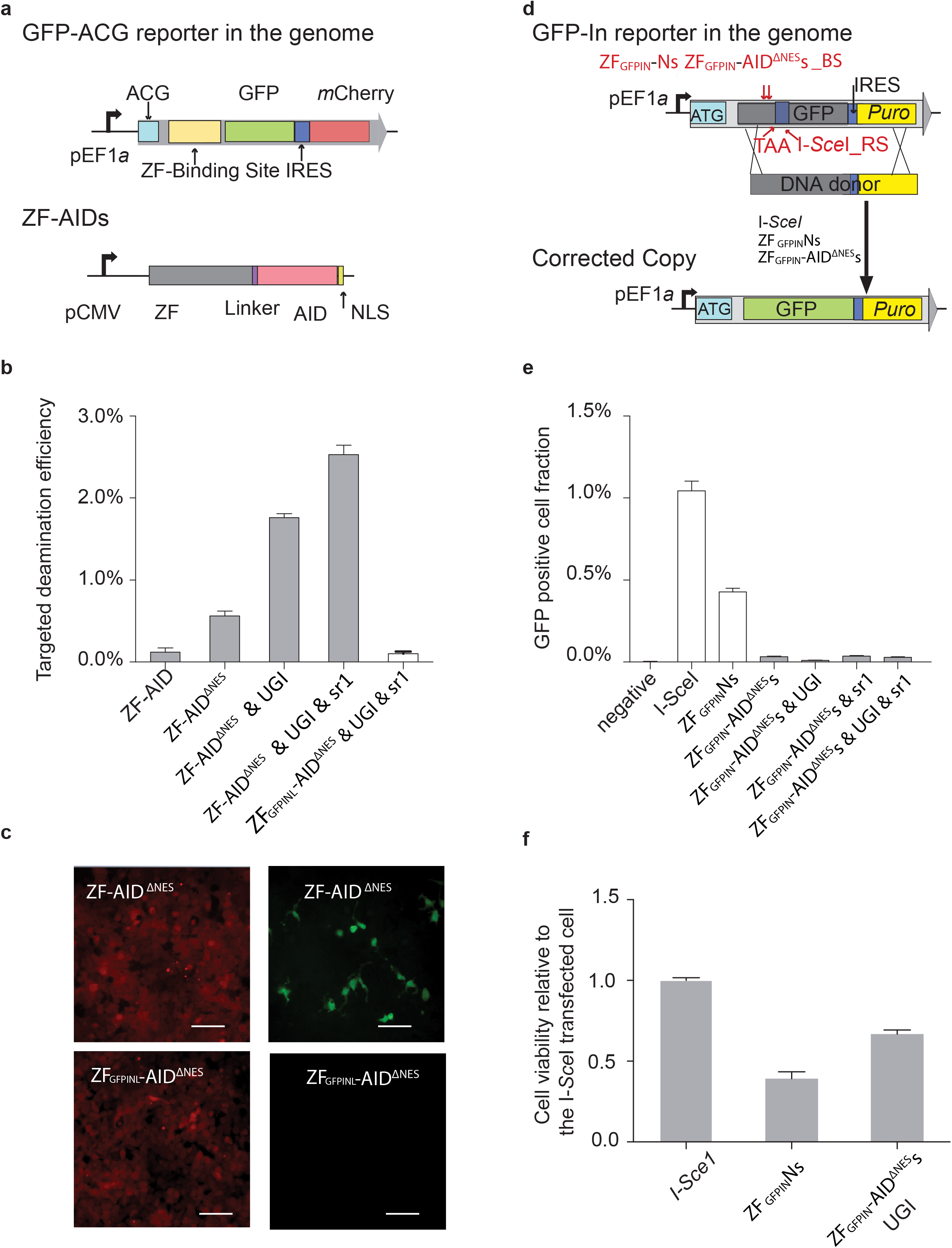
Targeted deamination and low toxicity of ZF-AID in human cells. **a**, Schematic representation of the ACG-GFP reporter system in HEK239FT cells (upper) and the ZF-AID (lower) tested for targeting deaminase activity. IRES, internal ribosome entry site; NLS, nuclear localization signal. **b**. Targeted deamination activity of ZF-AIDs. ACG-GFP reporter cells were transfected with the constructs labeled on the X-axis. Targeted deamination frequency was estimated as the proportion of GFP-rescued cells 48h after transfection. ZF-AID^ΔNES^ is identical to ZF-AID except with a deleted AID nuclear export signal (NES); UGI, inhibitor of UNG; sr1, shRNA-MSH2. c. ACG-GFP reporter cells imaged under fluorescence (mCherry (left)/GFP (right)) 48hr after transfection with ZF-AID ^ΔNES^ /UGI/sr1 or ZF _GFPINL_-AID^ΔNES^ /UGI/sr1 plasmids. Scale bar = 200 m **d**. Schematic design of DSBs assay. The genomically integrated GFP-In reporter includes a 35bp frame-shift insertion bearing a stop codon and I-SceI recognition site (I-SceI_RS). Of note, ZF_GFPIN_Ns and ZF_GFPIN_-AID^ΔNES^s binding sites (ZF_GFPIN_Ns/ZF_GFPIN_-AID^ΔNES^s_BS) were identical and located 82bp upstream of the insertion. We transfected the cells with a DNA donor carrying the wild-type GFP sequence along with I-Sce1/ ZF_GFPIN_Ns / ZF_GFPIN_-AID^ΔNES^s expression plasmids and assessed the DSB-generating rate by measuring HDR frequency as determined by GFP rescue of the cells. **e**. GFP rescue results determined by flow cytometry. Negative control was transfected with the DNA donor only. **f**. Cytotoxicity assay for ZF_GFPIN_-AID/UGI relative to I-Sce1. Detailed methods are in **Methods**. A value of <1 shows decreased cell survival as compared to I-S*ce*I, and demonstrates a toxic effect.

We next investigate toxicity of the targeted deaminase in human cells. To test whether ZF-AID^ΔNES^ can be safely used as a genome editing tool without incurring DSBs, we generated a HEK293FT reporter cell line carrying a non-functional frame-shifted GFP, which could be rescued by DSB-enhanced HDR with an exogenous donor DNA^3, 28^ (GFP-In reporter in Fig. 4d). If DSBs occurred from ZF-AID^ΔNES^ treatment, GFP+ cells can be detected. The GFP-In reporter also carries recognition sites for alternative nucleases, I-SceI and ZF_GFPIN_Ns (ZF_GFPINI_N & ZF_GFPINR_N)^28^. While expression of I-SceI and ZF_GFPIN_Ns generated 1.01% and 0.43% GFP+ cells respectively, expression of ZF_GFPIN_-AID^ΔNES^s gave rise to just 0.03% GFP+ cells, which was further reduced to 0.01% if the UNG inhibitor UGI was co-transfected^26, 29^ (Fig. 4e). Thus, targeted deaminase with UGI treatment created 40-fold fewer DSBs at the target locus than alternative nuclease-based approaches, close to negative control levels where only the DNA donor was delivered. The extent to which ZF-AID^ΔNES^ generated DSBs (0.01%) was very low compared to its C:G→T:A editing activity (2.1%, Fig. 4b). Furthermore, we observed higher cell survival in the ZF_GFPIN_-AID^ΔNES^s/UGI-expressing population (66%) than in the ZF_GFPIN_Ns-expressing population (41%, Fig. 4f), suggesting that targeted deaminases are less cytotoxic than ZFNs. Thus, expression of chimeric AID ^NES^s with UGI enables efficient genome editing in human cells without generating DSBs and with low cytotoxicity.

## Discussion

Our study demonstrates that fusing cytidine deaminases with DNA binding modules enables site-specific deamination of genomic loci in both prokaryotic and eukaryotic cells. We designed and optimized the structure of targeted deaminases to effectively convert a specific C:G base pair to T:A in the *E.coli* genome, achieving up to 13% editing frequency. We then applied the optimized chimeric deaminases to a human cell line and found that these novel enzymes could create site-specific single-nucleotide transitions in as many as 2.5% of cells. The transfected cells demonstrated decreased cytotoxicity compared with targeted nucleases. We found that inhibition of uracil and mismatch repair were critical to achieving these high editing rates.

A recent study independently developed targeted deaminases using Cas9 as the DNA binding domain and reported obtaining similar results, albeit by different means^30^. Their Cas9 deaminase achieved higher efficiencies by taking advantage of the fact that Cas9 binding generates an ssDNA loop, a natural substrate for AID and APOBEC deaminases. Instead of expressing UGI independently, they fused it to nikase-Cas9, and cleverly supressed mismatch repair by allowing Cas9 to nick the non-targeted strand.

The authors of this study suggested that, due to their efficiency and their avoidance of mutagenic dsDNA cuts, targeted deaminases might be promising tools for the correction of genetic diseases. However, our results suggest that further engineering is needed to reduce their processivity^30^ and off-target activity. First, it is still difficult to pinpoint the activity of targeted deaminases to a specific cytidine within a +/− 15bp window (Fig. 3c and 3d), and we also detected off-target mutations > 150bp away from the DNA binding site. These findings indicate that targeting does not eliminate the processivity of deaminases and suggest the need to engineer them to reduce off-targeting. Additionally, our whole genome sequencing data demonstrated elevated levels of global deamination in off-target WRC sites (Fig. 3e and 3f), suggesting that deaminases maintain their intrinsic DNA binding preferences and editing activity even when fused to targeted DNA binding domains. That the overexpression of APOBEC3B has been implicated as a driver for human breast, ovarian and cervical cancers through its generation of random C:G→T:A transitions^31, 32^ provides a cautionary note about the need to constrain the intrinsic preferences and activity of deaminases. One possible solution to this problem would be to create obligatory dimeric targeted deaminases by splitting the deaminase protein and fusing each half to an independent DNA binding domain, so that an active deaminase protein would only be generated if both halves are targeted to two specific nearby DNA sequences. This approach could also reduce the processivity of the enzyme.

Our results set the stage for the future engineering of additional targeted DNA nucleotide mutases beyond cytidine deaminases, such as targeted adenosine deaminases^33^, that effect changes to DNA without introducing DNA cuts or nicks. Aside from using such targeted mutases for gene therapy as suggested for deaminase in the recent study^30^, suitably engineered to reduce their intrinsic mutability as we indicate, we foresee other potential uses of these tools. First, as nucleases tend to be toxic in prokaryotes^34, 35^ due to their lack of efficient DSB repair pathways such as NHEJ, targeted DNA mutases may provide an effective means to engineer prokaryotic species such as *Streptococcus* for which few molecular tools are available. As demonstrated here with deaminases, targeted mutases have potential to be highly portable in both prokaryotic and eukaryotic systems. Second, although DSBs are better tolerated in eukaryotes, certain cell types are very sensitive to DSBs incurred by nucleases, such human induced pluripotent cells. We demonstrated that targeted deaminases incur significantly less cytotoxicity compared with nucleases in HEK293 cell and we envision that they and other targeted mutases will similarly be less toxic in these sensitive cell types. In addition, targeted mutases should be effective ways to generate precise mutations in non-dividing cells in which HDR activity is extremely low^13^. Moreover, the independency from exogenous DNA donor to make precise mutations likely allows this tool in the agriculture applications without GMO regulation, similar to recently successful cases of mushroom and corn in which CRISPR were used to disrupt endogenous genes. Finally, although our group demonstrated highly multiplexible (62X) targeting of a repetitive sequence in immortalized pig cells^36^, it has proven highly difficult to achieve this result in primary cells, likely because the high number of DSBs required to achieve this result leads to chromosomal rearrangements, senescence and apoptosis. We believe that the independence from DSBs may make targeted mutases a safer and more efficient tool for editing and studying repetitive elements in the genome.

## Methods

### Construction of fusion proteins

To construct ZFP-AID fusion proteins, we first PCR amplified ZFP from pUC57-ZFP^17^ and AID from pTrc99A-AID^37^ and fused these two parts with various linkers using overlap PCR. The fusion constructs were cloned into a pTrc-Kan plasmid. We fused AID with TALE by cloning AID into pLenti-EF1a-TALE(0.5 NI)-WPRE^20^ plasmid and then cloned TALE-AID fusions into the pTrc-Kan plasmid. APOBEC1, 3F, and 3G genes were synthesized (Genescript) and cloned into the pTrc-ZFP-Kan plasmid. To generate pCMV-ZF-AID constructs, we amplified ZF-AID cassette from pTrc-ZF-AID and cloned that into pCMV-hygo^20^ plasmid. The detailed construction methods are found in **Supplementary Method 1** and illustrated in **Supplementary Figure 4**. The sequences of the fusion proteins are listed in **Supplementary Sequence 1–5**.

### Construction of E.coli reporter cell lines

The GFP coding sequence was amplified from pRSET-EmGFP (Invitrogen). We modified the reporter by mutating the start codon to ACG and inserting a ZFP/TAL binding site upstream of the GFP coding sequence. To establish stable cell lines with a single copy of the GFP reporter sequence in the genome, we integrated the GFP cassette into the galK locus in the EcNR1 (MG1566 with λ-prophage::bioA/bioB) and EcNR2 (EcNR1 with mutS knocked out) strains^19^. To knock out ung, we replaced the ung gene with Zeocin resistance cassette via recombineering (**Supplementary Method 2**). In addition, all the reporter cell lines were transformed with pTac-T7polymerase to induce the expression of GFP. Subsequent modifications of the reporter were conducted using the MAGE system^19^. All sequences can be found in **Supplementary Sequence 6, 7**. Detailed protocols are available in **Supplementary Method 2**.

### E.coli cell culture and targeted deaminase activity assay

The reporter strains were electro-transformed with the plasmids coding for targeted deaminases. Single colonies were inoculated and cultured under 34°C in LB-min-media (5g NaCl, 5g yeast extract,10g tryptone in 1L ddH2O) supplemented with 100μg/mL Carbenicillin, 25μg/mL Chloramphenicol, 100μg/mL Spectinomycin, 100ug/mL Kanamycin. Targeted deaminase activities of the targeted deaminases were tested by inducing the expression the fusion protein with IPTG of final concentration 100μM when the O.D of the cell culture reached 0.4~0.6. To maintain the continuous cell proliferation, cell culture was diluted 100-fold into fresh media every 10 hours.

### Flow cytometry

Targeted deaminase activity as measured by GFP+ cell fraction in the total population was assayed by flow cytometry using a LSRFortessa cell analyzer (BD Biosciences). Bacteria culture was diluted 1:100 with PBS and vortexed for 30 seconds before flow cytometry. At least 100,000 events were analyzed for each sample. Targeted deamination efficiency was calculated as the percentage of GFP positive cells in the whole population.

### gfp gene Sanger sequencing

To genotype the GFP and GAPDH genes in *E.coli*, we inoculated single colonies in LB media and cultured them for 16 h at 34°C. PCR reactions with Phusion enzyme (NEB) were conducted with 1μl 100X diluted bacterial culture and Sanger sequencing were performed. Sequence of primers can be found in the **Supplementary Sequence 8** and the PCR protocol can be found in the **Supplementary Method 3**.

### Genomic library preparation

Corresponding reporter strains were transformed with ZFP-8aa-AID and TALE-C3-AID respectively. Single colonies were inoculated and split into the induction and non-induction groups. The expression of the deaminases was induced for 10 hours and the cell culture was plated on IPTG containing agar plate to isolate single colonies. After approximately 24h, we inoculated single colonies into LB-min media and cultured them overnight at 34°C. In order to extract chromosomal DNA and minimize the amount of plasmid DNA, Miniprep was first performed (Qiagen) according to manufacturer’s protocol and the sodium acetate/SDS precipitate formed was resuspended in the Lysis Buffer Type 2 (Illustra bacteria genomicPrep Mini Spin Kit, GE Healthcare) and the genomic DNA was recovered following manufacturer’s instructions (Illustra bacteria genomicPrep Mini Spin Kit, GE Healthcare). Genomic DNA libraries were constructed from 1.5–2 μg of genomic DNA. DNA was sheared in TE buffer (10 mM Tris (pH 8.0) 0.1 mM EDTA) using microTube (Covaris) with recommended protocol. Median DNA fragment sizes, estimated by gel-electrophoresis, were 150–250 bp. Sheared fragments were processed with the DNA Sample Prep Master Mix Set 1 (NEB). Adaptors consisted of the Illumina genomic DNA adaptor oligonucleotide sequences with the addition of 2-bp barcodes. Eight barcoded genomic libraries were pooled with equal molar amount. Detail protocols can be found in the **Supplementary Method 4**. The sequences of the adaptors and primers can be found in the **Supplementary Sequence 9**.

### Genomic DNA sequencing analysis

The reference genomic sequence for the reporter strain was generated by manually modifying the FASTA sequence of E. coli K-12 strain MG1655 to reflect the removal of *mutS* and *ung*, the insertion of the lambda prophage genome into the *bioAB* operon, and the insertion of the GFP reporter into the *galK* cassette. Genomic libraries were single-end sequenced using an Illumina Genome Analyzer, generating 100bp reads. The reads were first assigned to samples according to their 2-bp barcodes by exact matching and reads with fewer than 60 bases of high-quality sequence were discarded. Sorted reads were then aligned to the reference genomes using the Breseq package. Match lengths of at least 40 bases were required for alignment. In addition to Breseq’s single nucleotide variation (SNV) calling functionality, the SAMtools package^38^ was used on the resulting BAM file to corroborate short indels and single nucleotide variants. To validate the result of Breseq, MAQ was used as a second method to align the raw reads to the reference genomes and to call SNVs. FastQ files containing the sequencing reads were split based on the barcode, and trimmed using the FASTX-toolkit library. The resulting fastQ files were mapped to the reference genomes with MAQ^39^. Single nucleotide substitutions were considered valid when supported by a minimum read depth of 10 or a Phred-like consensus quality higher than 80. Finally, these three sets were merged to generate the final SNV set. SNVs called by both MAQ and SAMtools, or SNVs called by one and also called by Breseq, were kept. Indels were called by SAMtools alone. Breseq was used to identify new junctions using candidates generated by split read alignment. LiftOver^40^ was used to map the SNVs back to the original MG1655 genome (NCBI accession: NC_000913) for annotation. SNV effect prediction was done using the snpEff package^41^ and BioPerl. The analysis flow map (**Supplementary Fig.4**) provides information about the number of raw sequence reads, aligned reads, genome coverage and validated SNVs (**Supplementary Table 1 and 2**), and the list of SNVs (**Supplementary Table 3**) can be found in the supplementary information.

### Statistical analysis of the whole-genome sequence data

Wilcoxon test was used to analyze whether the mutation rate was higher in the strains with TALE-AID or ZF-AID induction. Intended mutations in TALE-AID, and ZF-AID strains were discarded for this analysis. X (SNVs not induced)=31, 21; Y(SNVs induced)=40, 31, 27, 33, 20, 32. H_0_: There is no difference in the mutation rate; H1: induced strains have a higher mutation rate. P_value_=0.25. The null hypothesis cannot be rejected. Therefore, there was no significant difference in the number of mutations between induced and uninduced strains. Due to the limited sample size, sensitivity simulations were performed to ensure an appropriate type II error. 1) Random samples from the observations of size m+n were taken, and divided in two groups A (n members) and B (m members). 2) An arbitrary value δ was added to B. 3) Wilcoxon one-sided p-value was calculated for comparison of groups A and B. A p-value under 0.05 was considered a success and recorded; otherwise a failure was recorded. 4) Steps 1 to 3 were repeated 10,000 times. The estimated power of the test was approximated by the proportion of successes among the 10,000 repetitions. 5) Steps 1 to 4 were repeated for a range of values of m+n and a range of δ values. The results are presented in the **Supplementary Figure 5**. With the current sample size, we could detect an increase of 13 SNVs, or higher (Statistical power=0.8).

### Poisson based modeling of number of genome edited sites

There are four sites in the genome with equivalent features as the targeted site. All of them contain an exact ZF binding sequence 11 bp away from an upstream WRC motif. Deamination was only detected in the targeted site with a maximum frequency of 7%. Assuming that alterations of these sites are Poisson distributed with *╭* = .07, the probability of detecting a second mutation in any strain is 0.03, and the probability *P* of *not detecting* an additional mutation in any of the 3 ZF-AID strains is 0.90.

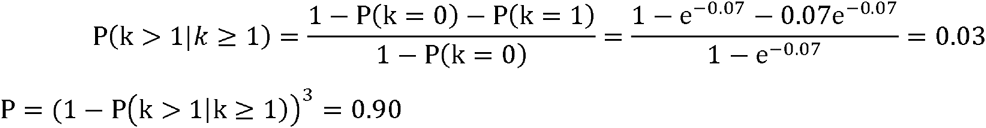

### Human cell culture

The human embryonic kidney cell line HEK293FT (Invitrogen) and the derivative reporter cell lines was maintained under 37 °C, 5% CO_2_ using Dulbecco’s modified Eagle’s Medium supplemented with 10% FBS, 2 mM GlutaMAX (Invitrogen), 100 U/ml penicillin and 100 μg/ml streptomycin.

### Targeted deaminase activity assay

The GFP-ACG reporter cell lines were generated by lentiviral transduction with low virus titration to make sure at most one copy of the reporter can be integrated into the genome. Single cells were isolated via FACS based on mCherry signal (Beckman Coulter MoFlo). Deaminase activity was tested by transfecting reporter cells with plasmids carrying ZF-AIDs. Briefly, HEK293FT cells were seeded into 12-well plates the day before transfection at densities of 4 × 10^5^ cells/well. Approximately 24 h after initial seeding, cells were transfected using Lipofectamine 2000 (Invitrogen) and 1.6ug DNA (400 ng of ZF-AID expression plasmid and/or 20 ng of UGI expression plasmid, and/or 20ng of ShRNA-MSH6 expression plasmid (Sigma), and pUC19(Invitrogen) plasmid to 1.6ug) per well. After 48 h, cells were trypsinized from their culturing plates and resuspended in 200 μl of media for flow cytometry analysis. At least 25,000 events were analyzed for each transfection sample. The flow cytometry data were analyzed using BD FACSDiva (BD Biosciences). The reporter and constructs sequences can be found in the **Supplementary Sequences 10, 11, 12**.

### Genotyping of human cell

To genotype the GFP target locus in HEK293 cells, we picked single GFP+ monocolonies and added each to 10ul 1X prepGEM buffer and enzyme (ZyGEM). After cell lysis according to the manufacturer’s instructions, the bulk product was added to a PCR reaction containing Platinum Taq polymerase (invitrogen). PCR products were cloned in pCR™4-TOPO (invitrogen) and capillary sequenced by Genewiz.

### DSB generating potential assay

The GFP-In reporter^28^ cell lines were generated by lentiviral transduction and successful reporter insertions were selected via puromycin selection. GFP-In reporter cells were plated in 12-well plates the day before transfection at densities of 4 × 105 cells/well transfected using Lipofectamine 2000 (Invitrogen) and 1.6ug DNA (400 ng of ZF-AID/ZF-nuclease/I-Sce1 expression plasmid and/or 20 ng of UGI expression plasmid, and/or 20ng of ShRNA-MSH6 expression plasmid, 1ug of DNA donor pUC19 (Invitrogen) plasmid to 1.6ug) per well. 72 hours after the transfection, cell were trypsinized and resuspended in 200 μl of media for flow cytometry analysis. At least 25,000 events were analyzed for each transfection sample. The flow cytometry data were analyzed using BD FACSDiva (BD Biosciences). The constructs sequence can be found in the **Supplementary Sequences 13**.

### Cytotoxicity assay

The assays were conducted as described before^42^. Briefly, HEK293FT cells were seeded in 12-well plates (4×10^5^□cells/well) and transfected after 24□h with 200□ng of deaminase/ nuclease expression plasmids, 10□ng of pmaxGFP (Lonzon), and pUC19 to 2□μg using calcium phosphate-mediated protocol. After 2 and 5 days, the fractions of GFP-positive cells were determined by flow cytometry (BD Biosciences). The survivability was calculated as the percentage of GFP-positive cell surviving at day 5 divided by the percentage of GFP-positive cells determined at day 2 after transfection. This ratio was normalized to the corresponding ratio after I-S*ce*I transfection, to yield the percentage survival as compared to I-S*ce*I.

